# RBD-based high affinity ACE2 antagonist limits SARS-CoV-2 replication in upper and lower airways

**DOI:** 10.1101/2023.06.09.544432

**Authors:** Matthew Gagne, Barbara J. Flynn, Christopher Cole Honeycutt, Dillon R. Flebbe, Shayne F. Andrew, Samantha J. Provost, Lauren McCormick, Alex Van Ry, Elizabeth McCarthy, John-Paul M. Todd, Saran Bao, I-Ting Teng, Shir Marciano, Yinon Rudich, Chunlin Li, Laurent Pessaint, Alan Dodson, Anthony Cook, Mark G. Lewis, Hanne Andersen, Jiří Zahradník, Martha C. Nason, Kathryn E. Foulds, Peter D. Kwong, Mario Roederer, Gideon Schreiber, Robert A. Seder, Daniel C. Douek

## Abstract

SARS-CoV-2 has the capacity to evolve mutations to escape vaccine-and infection-acquired immunity and antiviral drugs. A variant-agnostic therapeutic agent that protects against severe disease without putting selective pressure on the virus would thus be a valuable biomedical tool. Here, we challenged rhesus macaques with SARS-CoV-2 Delta and simultaneously treated them with aerosolized RBD-62, a protein developed through multiple rounds of *in vitro* evolution of SARS-CoV-2 RBD to acquire 1000-fold enhanced ACE2 binding affinity. RBD-62 treatment gave equivalent protection in upper and lower airways, a phenomenon not previously observed with clinically approved vaccines. Importantly, RBD-62 did not block the development of memory responses to Delta and did not elicit anti-drug immunity. These data provide proof-of-concept that RBD-62 can prevent severe disease from a highly virulent variant.

## Main Text

SARS-CoV-2 variants of concern (VOC) including B.1.351 (Beta), B.1.617.2 (Delta) and the currently circulating sublineages of B.1.1.529 (Omicron) have acquired mutations that enable substantial escape from neutralizing antibodies in convalescent or vaccinee sera (*1-7*). Efficacy against severe disease after two doses of mRNA COVID-19 vaccines has declined from ∼100% in clinical trials conducted at a time when ancestral strains were predominantly in circulation to 60-80% during the Omicron BA.1 wave (*8-12*), a result of both waning antibody titers and virus-acquired mutations. Boosting can restore protective efficacy but the benefit of boosting beyond a third dose is unclear (*13-15*) and accumulating evidence suggests that antigenic imprinting may offset the benefit of variant-matched boosts (*16-21*).

Antiviral therapeutic agents can reduce the effects of severe COVID-19 in individuals with or without prior immunity. Two drugs recently granted emergency use authorization by the FDA include Merck’s molnupiravir (Lagevrio) and Pfizer’s nirmatrelvir/ritonavir (Paxlovid). Interim data indicated that molnupiravir, a cytidine analog prodrug, reduced hospitalizations from COVID-19 by about 50% but further analysis has suggested a lower efficacy (*22*).

Nirmatrelvir/ritonavir has demonstrated substantial clinical efficacy, with an 89% decline in severe disease (*23*). However, nirmatrelvir/ritonavir is not routinely prescribed for the treatment of COVID-19 (*24, 25*). Furthermore, the emergence of drug-resistant mutations in the virus remains a possibility; while some have already been detected in people, no widely circulating variants currently demonstrate this capacity (*26-29*).

Consequently, there remains an urgent need for the development of additional therapeutic agents that reduce severe disease, particularly those that act in a variant-agnostic manner; that is, without directly targeting the virus. Host-targeted approaches could maintain their efficacy as new variants emerge even if they were so divergent from the ancestral strains such that antiviral drugs or prior immunity were rendered ineffective. We have previously described the development of an *in vitro* mutated SARS-CoV-2 receptor binding domain (RBD) that displays greatly enhanced binding to the virus target receptor, angiotensin converting enzyme 2 (ACE2), without inhibiting its natural enzymatic activity (Table S1). The product, termed RBD-62, has a binding affinity for ACE2 of 16 pM, an increase of 1000-fold compared to the wildtype Wuhan-Hu-1 (WT) RBD, which has a binding affinity of 1700 pM (*30*). *In vitro* models demonstrated its capacity to block infection of cell lines with a half-maximal inhibitory concentration (IC_50_) of 18 pM against the Beta variant. RBD-62 treatment of Syrian hamsters through inhalation at the time of infection with the ancestral strain USA-WA1/2020 (WA1) also resulted in protection against weight loss. Protection against severe disease in a model system more closely approximating humans or in the context of infection from a VOC that has been shown to result in substantial morbidity and mortality has previously not been established.

Here, we infected rhesus macaques with Delta which is the most pathogenic variant tested to-date in these animals. Macaques were treated immediately prior to challenge and every 24 hours for the next 5 days with RBD-62 administered to the airways via aerosolization. Protection was measured via titers of culturable virus and subgenomic virus RNA (sgRNA). We also analyzed mucosal and serum immune responses to Delta-specific antigens as well as to RBD-62 itself to assess any anti-drug antibody (ADA) responses.

## Results

### RBD-62 inhibits binding between the SARS-CoV-2 spike (S) and ACE2 in a variant-agnostic manner

Efficacies for COVID-19 therapies and vaccines have declined in the context of emerging variants, which has limited the long-term applicability of results established in initial clinical and pre-clinical research. To determine if data gathered on RBD-62 from a challenge model with an ancestral variant could be extrapolated to current and future emerging strains, we first used an *in vitro* assay to examine the ability of RBD-62 to block binding between ACE2 and S from a panel of different variants.

In contrast to unmutated WA1 RBD, which inhibited binding of both WA1 S and Delta S with a half-maximal inhibitory concentration (IC_50_) of ∼330 ng/mL, RBD-62 was nearly 100-times more potent with IC_50_ values of ∼4.5 ng/mL (Fig. 1A-B). Further, RBD-62 blocked binding between ACE2 and BA.1 S at almost the same concentration, and the IC_50_ for Beta S was only modestly higher at 6 ng/mL (Fig. 1C-D). Strikingly, IC_90_ values for RBD-62 against all variants were less than 40 ng/mL, whereas we were unable to achieve 90% binding inhibition for any variant using WA1 RBD as the inhibitor at any of our tested concentrations. As the potency of RBD-62 was strongly preserved across variants despite large differences in S sequence and binding affinity, we proceeded with *in vivo* evaluation of this product in a challenge model with Delta which, in our experience, is the most virulent variant and replicates at a high titer in the upper and lower airways of rhesus macaques.

**Fig. 1.**
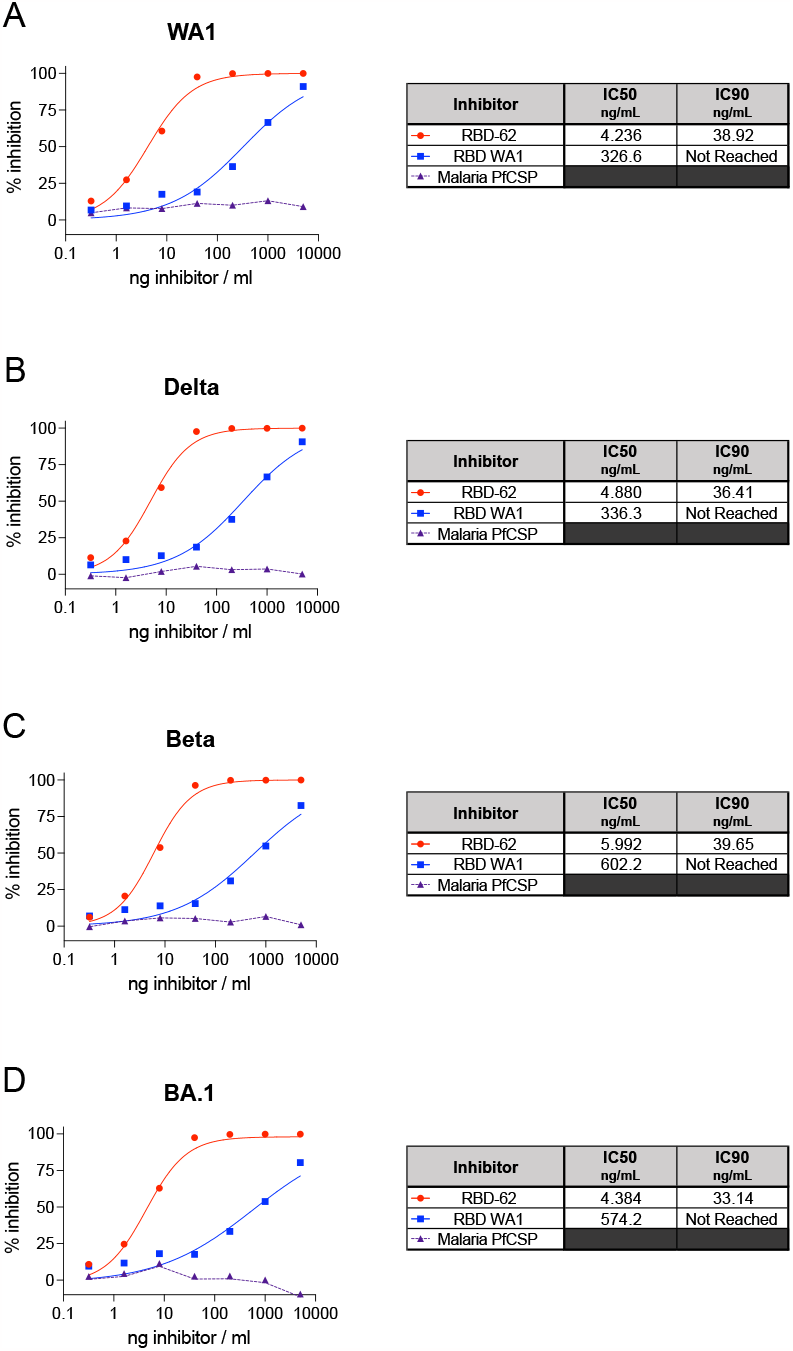
Inhibition of variant S-ACE2 binding. SARS-CoV-2 S from WA1 (**A**), Delta (**B**), Beta (**C**) and BA.1 (**D**) were mixed with soluble ACE2 in combination with indicated concentrations of RBD-62, RBD from WA1 or an irrelevant malaria protein (PfCSP) to determine percentage binding inhibition relative to maximum binding without inclusion of inhibitor. Icons represent average inhibition of duplicate technical replicates at each indicated dilution. IC_50_ and IC_90_ values (ng/mL) are indicated to the right of each graph and, along with the curves indicated on the graphs, were calculated using the nonlinear regression analysis tool in Prism. Dotted lines indicate background inhibition observed for PfCSP.

### RBD-62 protects rhesus macaques from Delta replication in the upper and lower airways

We delivered 2.5 mg RBD-62 to 8 rhesus macaques as an aerosol using the PARI eFlow® nebulizer as described previously (*31*) to target the drug to both the upper and lower airways. In addition, a further eight macaques received aerosolized PBS control. Both groups were challenged 1 hour later with Delta at a dose of 2×10^5^ TCID_50_. Primates continued to receive the same dose of aerosolized RBD-62 or PBS once per day for the next five days at which point treatment was stopped so that we could track the kinetics of virus rebound. Nasal swabs (NS) and bronchoalveolar lavage (BAL) were collected on days 2, 4, 7, 9 and 14 and RNA isolated for detection of virus replication by PCR for subgenomic RNA (sgRNA) encoding for the virus nucleocapsid (N) transcript (fig. S1).

We observed a significant decrease in virus sgRNA copies in the lungs on Day 2 with geometric mean of 2.3×10^5^ copies per mL BAL in the RBD-62 treatment group and 5.8×10^7^ copies in the PBS control group (*P*=0.0031). Likewise, RNA copies in the nose on day 2 were similar to the BAL, with geomeans of 2.8×10^5^ copies/swab in the treatment group and 4.5×10^7^ in the control group (*P*=0.0093) (Fig. 2A). Interestingly, the protective effect was no longer significant in either the nose or the lungs by day 7, which was the first collection timepoint after the cessation of RBD-62 treatment (*P*>0.05). Nonetheless, peak sgRNA copy numbers in the RBD-62 cohort were lower than in the control primates. For instance, while 4/8 control NHP had peak copy numbers greater than 10^8^ in either the lungs or the nose, virus titers never reached that level in any of the RBD-62-treated animals. Further, virus was cleared from the lower airway of all animals in the RBD-62 group by day 14 post-challenge whereas half of the control group still had detectable sgRNA at that timepoint.

**Fig. 2.**
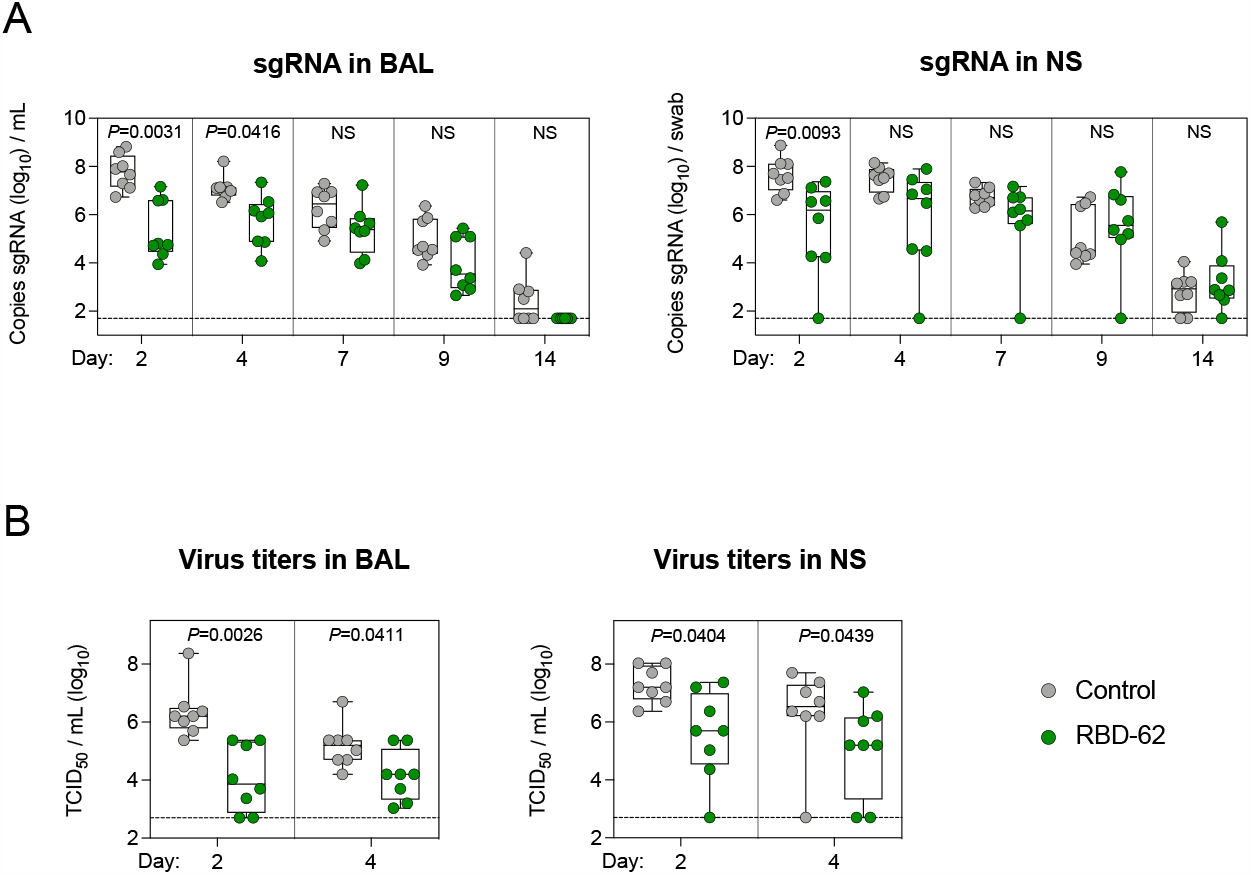
Delta replication in upper and lower airways. NHP (n=8 per group) were challenged with 2×10^5^ TCID_50_ Delta and simultaneously treated with RBD-62 (green circles) or PBS (gray circles). (**A**) Subgenomic RNA encoding for N transcript were measured in the upper and lower airways at days 2, 4, 7, 9 and 14 post-challenge. (**B**) Culturable virus was measured in the upper and lower airways at days 2 and 4 post-challenge. Dotted lines indicate assay limit of detection (LOD). Circles, boxes and horizontal lines represent individual animals, interquartile range and median, respectively. Statistical analyses shown for comparison of groups at each timepoint and were performed using Wilcoxon rank-sum test after a Holm’s adjustment across timepoints.

We also measured culturable virus in a tissue culture infectious dose assay (TCID_50_) which could indicate the potential for transmissibility (Fig. 2B). On day 2, the RBD-62 group had significantly less culturable virus than control animals, with geometric mean TCID_50_ values in the lungs of 1.1×10^4^ and 2.2×10^6^, respectively (*P*=0.0026). Culturable virus was also reduced in the upper airway, with TCID_50_ of 3.6×10^5^ for the RBD-62 group and 1.9×10^7^ for the control group (*P*=0.0404).

### Treatment with RBD-62 does not inhibit the induction of anti-SARS-CoV-2 humoral immunity

It is conceivable that the use of an ACE2-binding inhibitor during acute SARS-CoV-2 infection could prevent the formation of a primary or secondary immune response to the virus which would have been beneficial in the context of a future exposure. To test this hypothesis, we measured serum and mucosal IgG binding titers to a panel of variant RBDs, including WT, Delta and Omicron BA.1. At day 14 following challenge, titers to the Delta challenge stock were greater than either of the other strains for both the treated and untreated animals, indicative of a primary response. Geometric mean titers (GMT) to Delta rose from a baseline of 5×10^1^ to 2×10^9^ area under the curve (AUC) in the sera of the RBD-62 group by day 14 (Fig. 3A). While we observed a similar increase in GMT of control NHP by day 14, from 2×10^2^ to 2×10^10^ AUC, the kinetics were faster with evidence of a primary response as early as day 9. We next confirmed that there was a differential treatment effect across the entire 14-day time-course, which would indicate a blunted primary response due to a reduction in virus antigen in the RBD-62 group compared to the untreated group. Indeed, there was a significant treatment effect, with *P* = 0.0132. Likewise, mucosal binding titers to Delta RBD were higher on day 14 in the control primates, with GMT of 3×10^10^ in the lungs and 2×10^8^ in the nose as compared to 4×10^7^ and 2×10^7^ in the treated NHP respectively (*P* = 0.0001 in lungs and 0.0044 in nose) (Fig. 3B-C). Despite the attenuated response in the treated group, which reflected greater virus control by these animals, RBD-62 administration nevertheless did not preclude seroconversion.

**Fig. 3.**
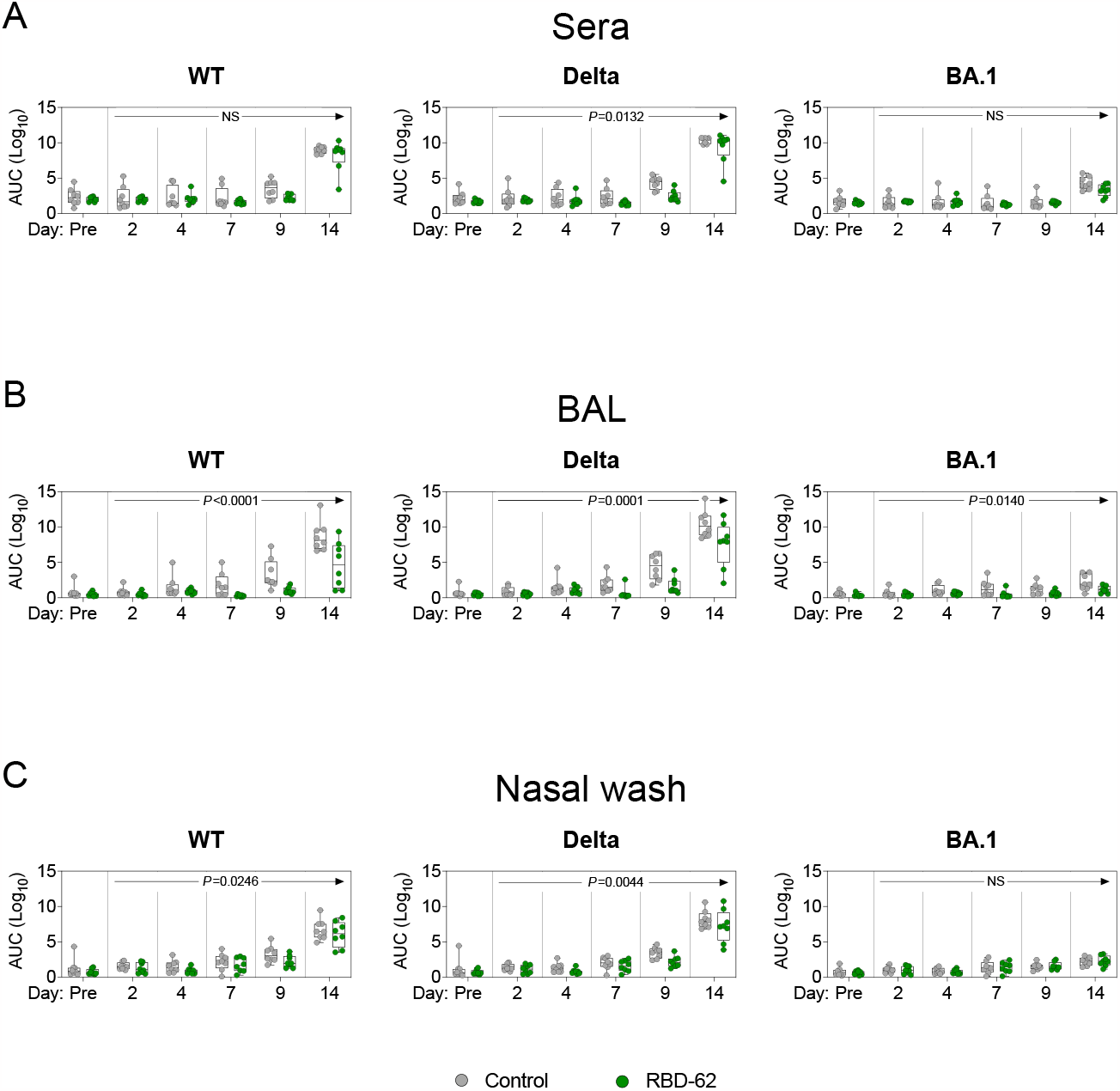
Anti-SARS-CoV-2 IgG binding titer kinetics. NHP (n=8 per group) were challenged with 2×10^5^ TCID_50_ Delta and simultaneously treated with RBD-62 (green circles) or PBS (gray circles). IgG binding titers were measured to wildtype, Delta and BA.1 RBD in the (**A**) sera, (**B**) BAL and (**C**) nasal wash one month prior to challenge (pre-challenge) and on days 2, 4, 7, 9 and 14 post-challenge. Sera were initially diluted 1:100 and then serially diluted 1:4. BAL and NW samples were initially diluted 1:5 and then serially diluted 1:5. Circles, boxes and horizontal lines represent individual animals, interquartile range and median, respectively. Statistical analyses shown for treatment effect (RBD-62 group vs. control group) across all post-challenge timepoints; longitudinal models were performed using generalized estimating equation.

Although treated animals developed antibodies to variant RBDs, it is possible that this response largely reflected or was enhanced by anti-drug immunity as RBD-62 is itself derived from WT RBD. To confirm that the immune response was not limited only to RBD, we quantified binding to various proteins and domains including RBD, whole S and nucleocapsid (N). Further, as we focused our analysis on WT protein, we reported responses using the WHO-determined standard antibody units for WT virus. We detected evidence of a primary response to RBD and S in the sera and BAL of control animals by day 9, while binding titers in the treated animals were not noticeably greater than background until day 14 (fig. S2a-b). Similarly, animals in the RBD-62-treated group mounted primary responses to WT nucleocapsid but with delayed kinetics and decreased magnitude as compared to the control cohort. On day 14, anti-nucleocapsid geometric mean titers (GMT) reached 11.4 antibody units per mL (BAU/mL) in the sera and 0.1 in the BAL of controls compared to 3.5 and 0.01 in the RBD-62 group. Titers in the nasal wash (NW) were markedly lower for both groups than in the BAL or sera (fig. S2c).

To further explore the effect of RBD-62 on the development of mucosal responses, we measured IgA binding titers to the aforementioned proteins (fig. S3). In agreement with our findings on IgG, the kinetics of the IgA response were faster in controls than in treated animals, with evidence of increased titers to RBD and S in the BAL of the controls by day 9 compared to day 14 in the RBD-62 group. However, we were nonetheless able to detect binding of IgA to both RBD and S in the BAL and NW of the RBD-62 group. Mucosal IgA responses to N were not clearly above background for either group of animals. Together with the data on mucosal IgG responses, this would suggest that a future mucosal vaccine boost would likely not be affected by prior RBD-62 treatment.

We have previously used the ACE2-RBD binding inhibition assay as a surrogate for neutralization in the mucosa (*16, 32, 33*). Here, we were able to detect inhibition for WT and Delta variants in the BAL of both control and treated groups (fig. S4a). We calculated that the median inhibitory capacity of BAL antibodies from the RBD-62 group was 6% against WT RBD and 8% against Delta RBD. Omicron BA.1 RBD-ACE2 binding inhibition was not detected, likely due to the divergence between the challenge stock and BA.1. NW inhibitory antibodies were not detectable in either group of animals (fig. S4b).

### Treatment with RBD-62 does not inhibit induction of anti-SARS-CoV-2 T cell responses

T cell epitopes present within SARS-CoV-2 are highly conserved (*16, 34*), suggesting that while newly emerging variants may continue to escape humoral immune responses, protection arising from T cell immunity may still be preserved. Thus, we next measured T cell responses to WT S peptides to determine if the administration of RBD-62 would interfere with their induction (fig. S5). Again, the kinetics of this response were faster in the control animals, with measurable increases in T_H_1 and CD40L^+^ T_FH_ responses in the periphery by day 7 compared to day 9 in the treated group (Fig. 4A, D). S-specific T_H_1 responses reached a median frequency of 0.2% in the controls and 0.1% in the treated NHP by day 14 although the difference was greater in the BAL (Fig. 4F). We did not detect T_H_2 responses in either the circulation or BAL (Fig. 4B, G). While we were able to detect some S-specific CD8^+^ T cells in the circulation of both groups, responses were much higher in the lungs — the primary site of virus replication — with median frequencies of 0.4% in both groups (Fig. 4C, H). We were also able to detect T cell responses to N in the circulation of the RBD-62 group (fig. S6).

**Fig. 4.**
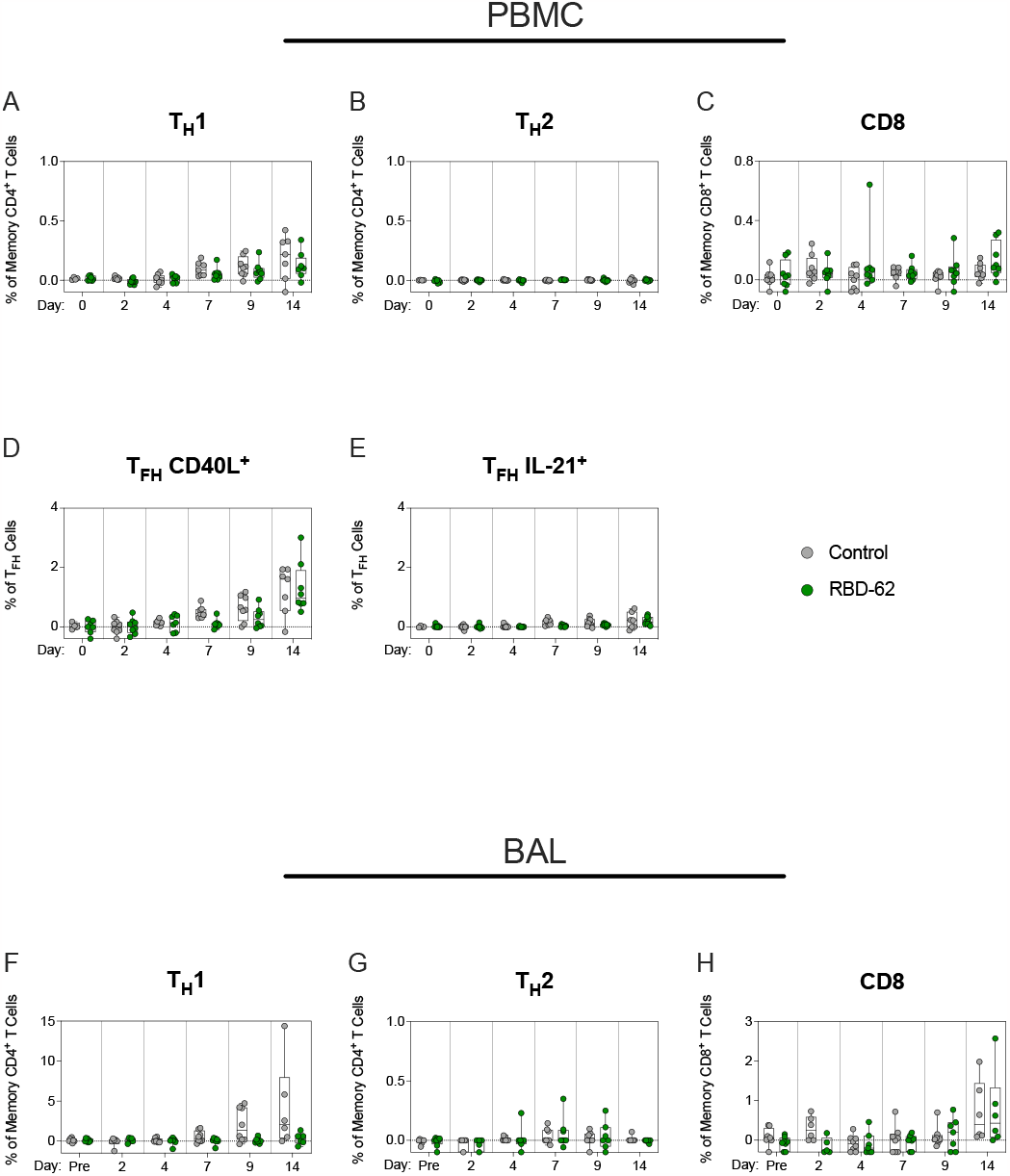
Kinetics of primary T cell responses following Delta challenge. NHP (n=8 per group) were challenged with 2×10^5^ TCID_50_ Delta and simultaneously treated with RBD-62 (green circles) or PBS (gray circles). (**A-E**) Peripheral blood mononuclear cells (PBMC) or (**F-H**) lymphocytes from BAL were collected prior to challenge (immediately preceding challenge for PBMC and one month pre-challenge for BAL) and on days 2, 4, 7, 9 and 14 post-challenge. Cells were stimulated with WA1 S1 and S2 peptide pools and responses measured by intracellular cytokine staining (ICS). (A and F) Percentage of memory CD4^+^ T cells expressing T_H_1 markers (IL-2, TNF or IFNγ). (B and G) Percentage of memory CD4^+^ T cells expressing T_H_2 markers (IL-4 or IL-13). (C and H) Percentage of memory CD8^+^ T cells expressing IL-2, TNF or IFNγ. (D and E) Percentage of T_FH_ cells expressing CD40L or IL-21, respectively. Dotted lines set at 0%. Reported percentages may be negative due to background subtraction and may extend beyond the lower range of the y-axis. Circles, boxes and horizontal lines represent individual animals, interquartile range and median, respectively.

### RBD-62 administration does not impair B cell memory to SARS-CoV-2 and does not elicit anti-drug immunity

Memory B cells are essential for mounting secondary responses upon boosting or reinfection and, together with long-lived plasma cells, form the basis of long-term humoral immunity (*35-38*). Due to imprinting, the initial exposure to virus establishes B cell antigen specificity and determines the capacity of the immune system to recognize novel variants (*16-21*). We therefore collected memory B cells from the peripheral circulation on day 14 following challenge and measured binding to pairs of fluorescently-labeled variant S-2P probes (fig. S7). Control and RBD-62-treated animals displayed almost identical patterns (Fig. 5A-B); out of all Delta and/or WA1-binding memory B cells in the controls, a geometric mean frequency of 42% bound to Delta alone compared to 49% in the treated animals. In the controls, 55% of the population was cross-reactive and capable of binding both S compared to 46% in the RBD-62 group. When examining the pool of memory B cells capable of binding to Delta and/or BA.1, only 37% and 35% were dual-specific in the controls and the RBD-62 group, respectively. This smaller fraction of BA.1/Delta-specific cross-reactive cells as compared to the WA1/Delta-specific cross-reactive pool is likely due to the reduced number of shared epitopes between BA.1 and Delta. We also measured the frequency of S-specificity among all memory B cells (fig. S8). As most recently circulating strains of SARS-CoV-2 are within the Omicron lineage, any limitation that RBD-62 treatment would place on the development of BA.1-binding B cell responses would be especially concerning. However, the frequencies of BA.1/Delta-specific cross-reactive memory B cells were similar between the two cohorts with geometric mean frequencies of 0.14% in the controls and 0.13% in the treatment group (fig. S8b).

**Fig. 5.**
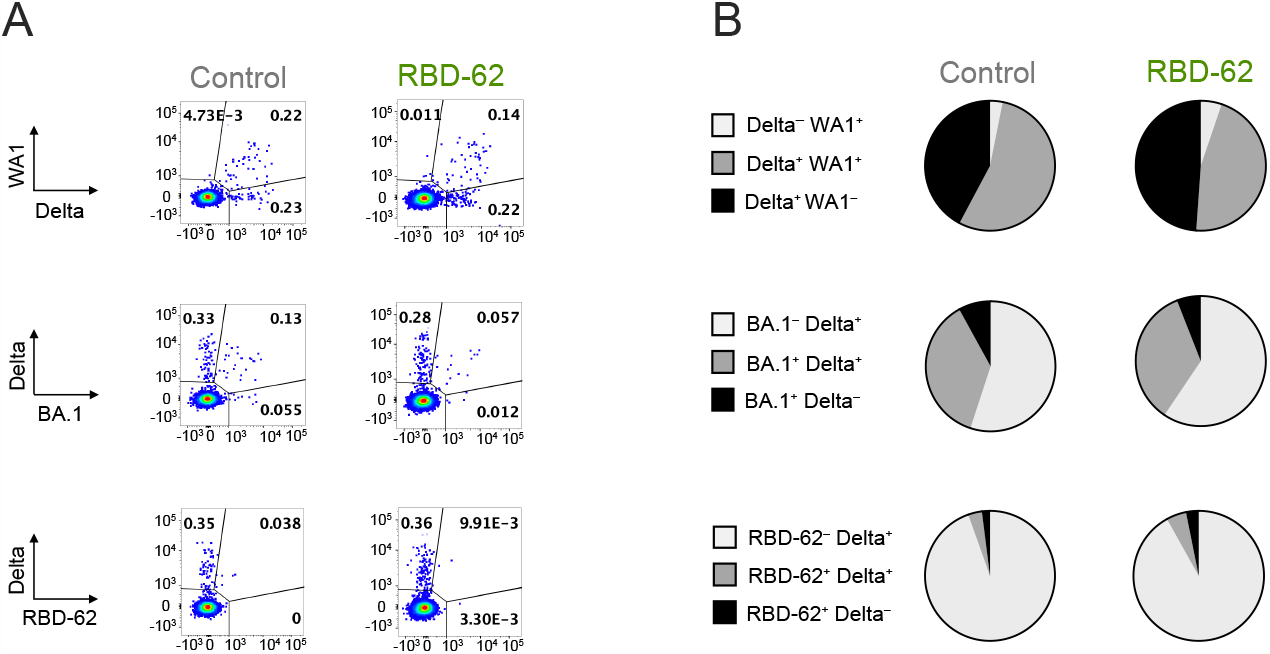
Memory B cell responses following Delta challenge and RBD-62 treatment. NHP (n=8 per group) were challenged with 2×10^5^ TCID_50_ Delta and simultaneously treated with RBD-62 or PBS. Memory B cell specificity was determined at day 14 post-challenge via binding to fluorochrome-labeled variant probe pairs as indicated in figure legends. Probe pairs include WA1 and Delta S-2P, Delta and BA.1 S-2P, and Delta S-2P and RBD-62. (**A**) Representative flow cytometry graphs for one animal in the control group (left) or treated group (right). Event frequencies denote proportion of probe-binding cells within the total class-switched memory B cell population. Cross-reactive memory B cells are represented by events in the top right quadrant whereas single-positive memory B cells reside in the top left or bottom right quadrants. (**B**) Pie charts indicating the geometric mean frequency of the entire S-specific memory B cell compartment capable of binding to both members of a variant probe pair (dark gray) or a single variant within the pair (light gray or black). Control group displayed on left and treated group displayed on right.

We next expanded our analysis of memory B cell binding specificities to include recognition of RBD-62. Two weeks following challenge and initiation of RBD-62 treatment, only 8% of memory B cells in the treatment group with specificities for RBD-62 and/or Delta bound to RBD-62. This was not meaningfully different from the control group (Fig. 5). Further, as a frequency of total memory B cells, the RBD-62-binding population following treatment was hard to distinguish from the pre-challenge background staining (fig. S8c).

## Discussion

As newly emerging SARS-CoV-2 variants continue to evolve while evading prior immunity, acquiring mutations which could render existing anti-viral drugs ineffective and even potentially increasing their affinity for ACE2 (*2-7, 27-29, 39-42*), it is essential to develop new treatments for COVID-19 which are variant-agnostic. However, to the best of our knowledge, no host-targeting treatments that are designed specifically to reduce virus load have been tested in clinical trials or within an advanced preclinical challenge model such as nonhuman primates. Here we describe the *in vivo* validation in rhesus macaques of RBD-62, a therapeutic agent which was designed through *in vitro* evolution to outcompete SARS-CoV-2 RBD for binding to human ACE2 while not interfering with the receptor’s natural enzymatic activity. Targeting the host receptor rather than the virus itself has several benefits including the avoidance of selective pressure and the ability to retain efficacy despite the emergence of new variants, which are unlikely to achieve the 1000-fold increase in ACE2-binding affinity needed to gain a competitive advantage over RBD-62. Moreover, optimizing aerosol delivery of this protein (*31*) promotes specific targeting of the respiratory track, including the lungs, achieving a significant reduction in virus replication for the duration of treatment. Importantly, after drug delivery was terminated, virus titers remained lower in the BAL. At the therapeutic dosage used here, RBD-62 did not inhibit the induction of serum or mucosal IgG or IgA responses to the challenge virus or other variants tested. T cell and B cell immunity were also preserved with no evidence of anti-drug immunity which would have precluded the reuse of this therapeutic agent upon subsequent reinfection.

The benefits of an ACE2 antagonist could theoretically extend to other host-targeting approaches including anti-ACE2 antibodies (*43, 44*) and ACE2 decoys that bind to RBD (*45-55*). It is noteworthy that chiropteran ACE2 functions as a host receptor for NeoCoV and PDF-2180, two merbecoviruses which have so far been restricted to transmission within bats (*56*). Continued work on developing biomolecules such as RBD-62 that can block the interaction between SARS-CoV-2 S and human ACE2 has potential benefits not only in this current pandemic but also in our continued vigilance against potential spillover events.

Advancement of this therapeutic agent, or similar host-targeting drugs, into clinical trials would require further optimization of dosage. It is possible that higher doses of RBD-62, or a longer duration of treatment, would have further suppressed virus replication, maintaining a significant protective effect until complete clearance. However, any advantage provided by increasing the amount of RBD-62 would have to be balanced with the possibility of inducing anti-drug antibody responses.

It has not escaped our notice that the loss of protection following treatment cessation coincided with a delayed primary immune response as indicated by slower kinetics of measurable IgG and IgA titers as well as ACE2-binding inhibitory antibodies and T cell responses. This is likely due to the low levels of virus antigen resulting from RBD-62 treatment, a conclusion supported by a recent publication on the muted primary response arising during nirmatrelvir treatment (*57*). This would suggest that an increase in the amount of antigen available to elicit an immune response without a commensurate increase in virus replication may be beneficial during RBD-62 treatment. Thus, one approach that may be worth exploring would be vaccination at the time of treatment.

Additionally, the protective effect of RBD-62 was striking in that it was observed in both the nose and the lungs. Indeed, our previous findings using mRNA vaccines delivered intramuscularly to nonhuman primates have shown that protection is often either delayed or absent in the upper airway, likely due to the higher threshold of antibodies required for virus suppression in the nose as compared to the lungs (*16, 32, 33, 58, 59*). Aerosolization of RBD-62 through the PARI nebulizer enables efficient delivery to both the lungs and the nose, highlighting the potential for this medication not only to be administered in a hospital setting to reduce the severity of disease but also to block infection. Indeed, RBD-62 could be employed as a preventative agent for healthcare workers or immunocompromised individuals who cohabitate with an infected individual. In the context of widespread immunity from prior infections and the global vaccination campaign, any impact that the use of RBD-62 could have on transmission reduction could accelerate the transition of the current pandemic into an endemic phase.

Furthermore, the finding that RBD-62 maintains its potency against a panel of different variants stands in stark contrast to the declining efficacy of currently approved vaccines and previously authorized monoclonal antibodies. Indeed, there is no clinically available monoclonal antibody with the capacity to neutralize the currently circulating Omicron sublineages (*60-64*). Vaccine development is predicated on predictions of which strains will be dominant at a future time. This can result in both significant lag times between identification of new strains with transmission advantages and authorization of variant-matched vaccine boosts and also in a mismatch between the vaccine immunogen and the currently circulating variant. As an alternative approach, we have described a rationally designed therapeutic agent which can be used to treat or prevent COVID-19 regardless of future SARS-CoV-2 evolution.

## Acknowledgments

We would like to thank Ruth Woodward and the entire Translational Research Program, Vaccine Research Center, National Institute of Allergy and Infectious Diseases, National Institutes of Health for expert technical support regarding animal procedures. We would also like to acknowledge Matthew Burnett for assistance in figure design. Patricia Darrah provided guidance on operation of the PARI nebulizer. Lawrence Wang provided the malaria circumsporozoite protein which was used as a negative control in our ACE2 binding inhibition assay. We would also like to thank Josue Marquez and Anna Mychalowych for sample processing and handling.

## Author contributions

Conceptualization: MG, JZ, GS, RAS, DCD Methodology: MG, J-PMT, SM, YR, CL, JZ, GS, DCD

Investigation: MG, BJF, CCH, DRF, SFA, SJP, LM, AVR, EM, J-PMT, SB, I-TT, SM, YR, CL, LP, AC, JZ, MCN, GS

Visualization: MG, SFA, SJP, KEF, DCD

Project administration: MG, BJF, J-PMT, KEF, DCD

Supervision: MG, SFA, J-PMT, YR, LP, AD, MGL, HA, KEF, PDK, MR, GS, RAS, DCD

Writing – original draft: MG, DCD

Writing – review & editing: MG, BJF, CCH, DRF, SFA, SJP, LM, AVR, EM, J-PMT, SB, I-TT, SM, YR, CL, LP, AD, AC, MGL, HA, JZ, MCN, KEF, PDK, MR, GS, RAS, DCD

## Data and materials availability

All data are available in the main text or the supplementary materials.

