## Supplemental Files (Methods & Supplemental Figures) for "RBD-based high affinity ACE2 antagonist limits SARS-CoV-2 replication in upper and lower airways"

**The PDF file includes:**

Materials and Methods
Figs. S1 to S8
Table S1

### Materials and Methods

Experimental design

Four- to seven-year-old male Indian-origin rhesus macaques were sorted into two groups of 8 NHP based on age and weight. RBD-62 formulated in gelatin was administered to one group at the time of Delta challenge while PBS formulated in gelatin was administered to the other group. Animals were challenged with $2x10^5$ $TCID_{50}$ of Delta (BEI, NR-56116). $1.5x10^5$ $TCID_{50}$ was administered via intratracheal route, and $0.25x10^5$ $TCID_{50}$ was administered intranasally to each nostril. Animals were housed at Bioqual, Inc. and all procedures approved and conducted in accordance with regulations of Animal Care and Use Committees of Bioqual and the Vaccine Research Center of the National Institutes of Health.

Production and purification of RBD-62

The RBD-62 protein was produced in several batches to a total of 4.6L Expi293F cells (ThermoFisher) by transient transfection of the pCAGGS plasmid (*30*). Plasmid DNA purified by NucleoBond Xtra Midi kit (Macherey-Nagel) was transfected using ExpiFectamine 293 Transfection Kit (ThermoFisher) according to the manufacturer's protocol. Media was collected 72 – 96 hours post-transfection, when the cell viability decreased to 50%, by centrifugation (1500 rpm for 15 minutes). The media was clarified by filtration through a 0.45 µm Nalgene (ThermoFisher) filter and loaded on two 5 ml HisTrap Fast Flow columns (Cytivia) connected in series using ÄKTA pure system (Cytivia). The column was washed with 5 column volumes (CV) of PBS 20mM imidazole pH 7.4 and eluted by step elution of 60% elution buffer PBS, 500mM imidazole pH 7.4. Eluted protein was concentrated by Amicon Ultra Centrifugal Filter Units, MWCO 3 kDa (Merck Millipore Ltd), and uploaded onto Superdex 200 16/600 (Cytiva) preequilibrated in PBS. The purity and quality of the eluted protein were analyzed by SDS-PAGE and Tycho (Nanotemper), respectively.

RBD-62 inhibition of S-ACE2 binding

Inhibitors including RBD-62, WA1 RBD (VRC, NIH) and truncated *Plasmodium falciparum* circumsporozoite protein 5/3_SAmut (Robert Seder, VRC, NIH) were chosen for comparison due to similar molecular weight (~25 to 27 kDa). 5/3_Samut (*65*) sequence is available on GenBank (ID: MT891178.1). All proteins were diluted to 5 µg/mL and then serially diluted 5-fold. ACE2 binding inhibition assay was performed with V-Plex SARS-CoV-2 Panel 23 (ACE2) Kit (MSD) per manufacturer's instructions. Plates were read on MSD Sector S 600 instrument. All samples run in duplicate and normalized to the average luminescent units measured for each variant without the addition of inhibitor, with the average inhibition at each dilution indicated by the icons. $IC_{50}$ values (ng/mL) were calculated via the [Agonist] vs. normalized response - variable slope equation within the nonlinear regression analysis tool in Prism. $IC_{90}$ values (ng/mL) were calculated via the [Agonist] vs. response - Find ECanything within the nonlinear regression analysis tool in Prism with bottom and top y-values constrained to 0 and 100, respectively. $IC_{50}$ and $IC_{90}$ values not listed for irrelevant malaria protein.

RBD-62 administration

RBD-62 was provided by Gideon Schreiber (Weizmann Institute of Science) and formulated with gelatin as delivery vehicle. Gelatin (Sigma-Aldrich, G1890) was prepared at a concentration of 4 mg/mL in Dulbecco's PBS (Gibco). RBD-62 or PBS control was then mixed with gelatin at a 1:1 ratio. Each animal was administered 2.5 mg RBD-62 or PBS control at an effective concentration of 2 mg/mL gelatin with a total volume of 4.6 mL via pediatric mask attached to a Pari eFlow nebulizer (PARI GmbH) that delivered 4 µm particles deep into the lung of an anesthetized macaques, as previously described (*66*).

Subgenomic RNA quantification

sgRNA was isolated and quantified by researchers blinded to vaccine status as previously described (*58*). Briefly, total RNA was extracted from BAL fluid and nasal swabs using RNAzol BD column kit (Molecular Research Center). PCR reactions were conducted with TaqMan Fast Virus 1-Step Master Mix (Applied Biosystems), forward primer in the 5' leader region and N gene-specific probe and reverse primer as previously described (*16*):

sgLeadSARSCoV2_F: 5'-CGATCTCTTGTAGATCTGTTCTC-3'

N2_P: 5'-FAM- CGATCAAAACAACGTCGGCCCC-BHQ1-3'

wtN_R: 5'-GGTGAACCAAGACGCAGTAT-3'

Amplifications were performed with a QuantStudio 6 Pro Real-Time PCR System (Applied Biosystems). The assay lower LOD was 50 copies per reaction.

$TCID_{50}$ quantification of SARS-CoV-2 from BAL and NS

$TCID_{50}$ assay was conducted as described previously (*58*). Briefly, Vero-TMPRSS2 cells (VRC/NIH) were plated at 25,000 cells / well in Dulbecco's Modified Eagle Medium (DMEM) + 10% FBS + Gentamicin and the cultures were incubated at 37°C, 5.0% $CO_2$. Cells reached 80–100% confluence the following day. Medium was aspirated and replaced with 180 µL of DMEM + 2% FBS + gentamicin. Twenty µL of BAL or nasal swab sample was added to top row in quadruplicate and mixed using a P200 pipettor 5 times. Using the pipettor, 20 µL was transferred to the next row, and repeated down the plate (columns A-H) representing 10-fold dilutions. The tips were disposed for each row and repeated until the last row. Positive (virus stock of known infectious titer in the assay) and negative (medium only) control wells were included in each assay set-up. The plates were incubated at 37°C, 5.0% $CO_2$ for 4 days. The cell monolayers were then visually inspected for cytopathic effect (CPE). The $TCID_{50}$ value was calculated using the Read-Muench formula.

Serum and mucosal antibody titers

Quantification of antibodies in the blood and mucosa were performed as previously described (*67*). Briefly, total IgG and IgA antigen-specific antibodies to SARS-CoV-2-derived antigens were determined in a multiplex serology assay by Meso Scale Discovery (MSD). We measured responses using V-Plex SARS-CoV-2 Panel 22 for variant RBD and Panel 1 for WT proteins and protein domains according to manufacturer's instructions, except 25µl of sample and detection antibody were used per well. BAL and NW were initially concentrated 10-fold using Amicon

Ultra centrifugal filter 10kDa MWCO (Millipore). For measurement of antibody titers to variant RBD, concentrated BAL and NW were initially diluted 1:5 and then serially diluted 1:5; heat inactivated plasma was initially diluted 1:100 and then serially diluted 1:4. Data presented as AUC. For measurement of antibody titers to SARS-CoV-2 protein domains, concentrated BAL and NW were diluted at 1:5, 1:10 and 1:20 ratios; heat inactivated plasma was diluted at 1:25, 1:50 and 1:100 ratios. Results reported as BAU/mL based upon reference standard included in Panel 1 kit according to manufacturer's instructions.

RBD-ACE2 binding inhibition

BAL fluid and NW were concentrated 10-fold with Amicon Ultra centrifugal filter 10kDa MWCO (Millipore). To remove residual RBD-62 prior to binding inhibition assay, fluid was diluted 1:1 in 50mM sodium phosphate, 300mM sodium chloride (binding buffer). HisPur Ni-NTA spin plate (Thermo Scientific) was equilibrated with binding buffer and the diluted fluid was applied to the plate and incubated with agitation at 4C overnight. The purified fluid was collected after centrifugation for 1 minute at 1000 x *g*. The fluid was then dialyzed against Diluent 100 using Pierce Microdialysis Plates (Thermo Scientific). Purified fluid was diluted to a final ratio of 1:5. ACE2 binding inhibition assay was performed with V-Plex SARS-CoV-2 Panel 22 (ACE2) Kit (MSD) per manufacturer's instructions. Plates were read on MSD Sector S 600 instrument. Results are reported as percent inhibition.

Intracellular Cytokine Staining (ICS)

Cryopreserved PBMC and BAL cells were thawed and rested overnight in a 37C, 5% $CO_2$ incubator. The next morning, cells were stimulated with SARS-CoV-2 spike protein (S1 and S2, matched to ancestral COVID-19 mRNA vaccine insert) and nucleoprotein (N) peptide pools (JPT Peptides) at a final concentration of 2 μg/ml in the presence of 3 mM monensin for 6 hours. The S1, S2 and N peptide pools are comprised of 158, 157 and 102 individual peptides, respectively, as 15mers overlapping by 11 aa in 100% DMSO. Negative controls received an equal concentration of DMSO instead of peptides (final concentration of 0.5%). ICS was performed as previously described (*33, 68*). The following monoclonal antibodies were used: CD3 APC-Cy7 (clone SP34.2, BD Biosciences), CD4 PE-Cy5.5 (clone SK3, Thermo Fisher), CD8 BV570 (clone RPA-T8, BioLegend), CD45RA PE-Cy5 (clone 5H9, BD Biosciences), CCR7 BV650 (clone G043H7, BioLegend), CXCR5 PE (clone MU5UBEE, Thermo Fisher), CXCR3 BV711 (clone 1C6/CXCR3, BD Biosciences), PD-1 BUV737 (clone EH12.1, BD Biosciences), ICOS Pe-Cy7 (clone C398.4A, BioLegend), CD69 ECD (cloneTP1.55.3, Beckman Coulter), IFN-g Ax700 (clone B27, BioLegend), IL-2 BV750 (clone MQ1-17H12, BD Biosciences), IL-4 BB700 (clone MP4-25D2, BD Biosciences), TNF-FITC (clone Mab11, BD Biosciences), IL-13 BV421 (clone JES10-5A2, BD Biosciences), IL-17 BV605 (clone BL168, BioLegend), IL-21 Ax647 (clone 3A3-N2.1, BD Biosciences) and CD154 BV785 (clone 24-31, BioLegend). Aqua live/dead fixable dead cell stain kit (Thermo Fisher Scientific) was used to exclude dead cells. All antibodies were previously titrated to determine the optimal concentration. Samples were acquired on a BD FACSymphony flow cytometer and analyzed using FlowJo version 10.8.2 (Treestar, Inc., Ashland, OR).

B cell probe binding

Flow cytometric analysis of antigen-specific memory B cell frequencies was performed as previously described (*33*). Briefly, cryopreserved PBMC were thawed and stained with the following antibodies (monoclonal unless indicated): IgD FITC (goat polyclonal, Southern Biotech), IgM PerCP-Cy5.5 (clone G20-127, BD Biosciences), IgA Dylight 405 (goat polyclonal, Jackson Immunoresearch Inc), CD20 BV570 (clone 2H7, Biolegend), CD27 BV650 (clone O323, Biolegend), CD14 BV785 (clone M5E2, Biolegend), CD16 BUV496 (clone 3G8, BD Biosciences), CD4 BUV737 (clone SK3, BD Biosciences), CD19 APC (clone J3-119, Beckman), IgG Alexa 700 (clone G18-145, BD Biosciences), CD3 APC-Cy7 (clone SP34-2, BD Biosciences), CD38 PE (clone OKT10, Caprico Biotechnologies), CD21 PE-Cy5 (clone B-ly4, BD Biosciences) and CXCR5 PE-Cy7 (clone MU5UBEE, Thermo Fisher Scientific). Stained cells were then incubated with streptavidin-BV605 (BD Biosciences) labeled Delta S-2P, BA1 S-2P or RBD-62 and streptavidin-BUV661 (BD Biosciences) labeled WA1 or Delta S-2P for 30 minutes at 4°C (protected from light). Cells were washed and fixed in 0.5% formaldehyde (Tousimis Research Corp) prior to data acquisition. Aqua live/dead fixable dead cell stain kit (Thermo Fisher Scientific) was used to exclude dead cells. All antibodies were previously titrated to determine the optimal concentration. Samples were acquired on a BD FACSymphony cytometer and analyzed using FlowJo version 10.7.2 (BD, Ashland, OR).

Quantification and statistical analysis

Comparisons of animals that received RBD-62 vs control for virus titers and humoral responses post-challenge are based on Wilcoxon tests on individual days while longitudinal analyses are based on Generalized Estimating Equations (GEE). We adjusted for multiple comparisons across timepoints for each assay using Holm's adjustment; we did not adjust to account for multiple comparisons across different assays or different target antigens. *P*-values are shown in the figures, and the relevant statistical analyses and sample *n* are listed in corresponding figure legends. NS denotes that the indicated comparison was not significant, with $P>0.05$.

Virus titers were analyzed on the $\log_{10}$ scale; humoral responses were analyzed as the area under the curve (AUC) on the $\log_{10}$ scale. Values below the limit of detection or lower limit of quantification for virus titers were set to half of the value for statistical analysis (25 copies sgRNA or 1.35 $TCID_{50}$ per mL or per swab). Antibody binding and virus assays are log-transformed as appropriate. All statistical analyses were done using R version 4.2.1. All flow cytometry data graphed in FlowJo version 10.7.2 (B cell binding) or version 10.8.2 (ICS) while all other graphs designed using Prism version 9.3.1.

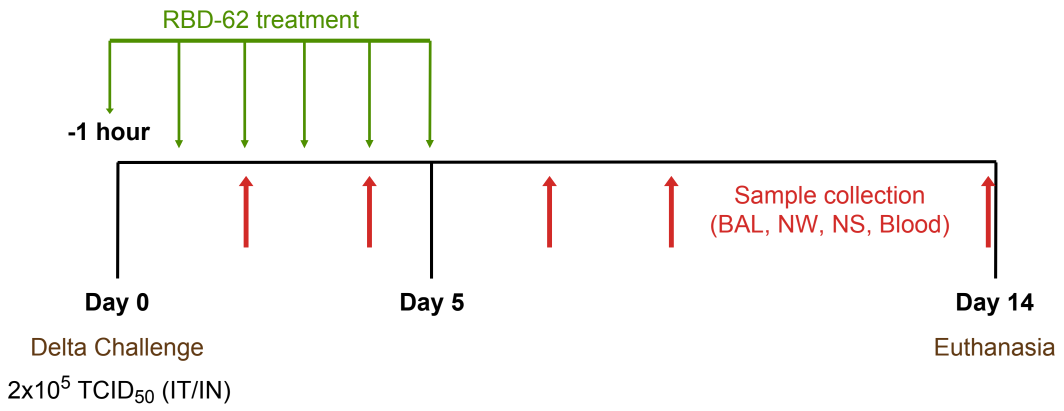


**Fig. S1.**
**Experimental schema.** NHP (*n*=8 per group) were challenged with $2x10^5$ $TCID_{50}$ Delta on day 0. RBD-62 was administered to one group of NHP on day 0 (1 hour prior to challenge) and on each successive day up to and including day 5. BAL, NW, NS and blood (both sera and PBMC) were collected at timepoints indicated by red arrows. All animals were euthanized on day 14.

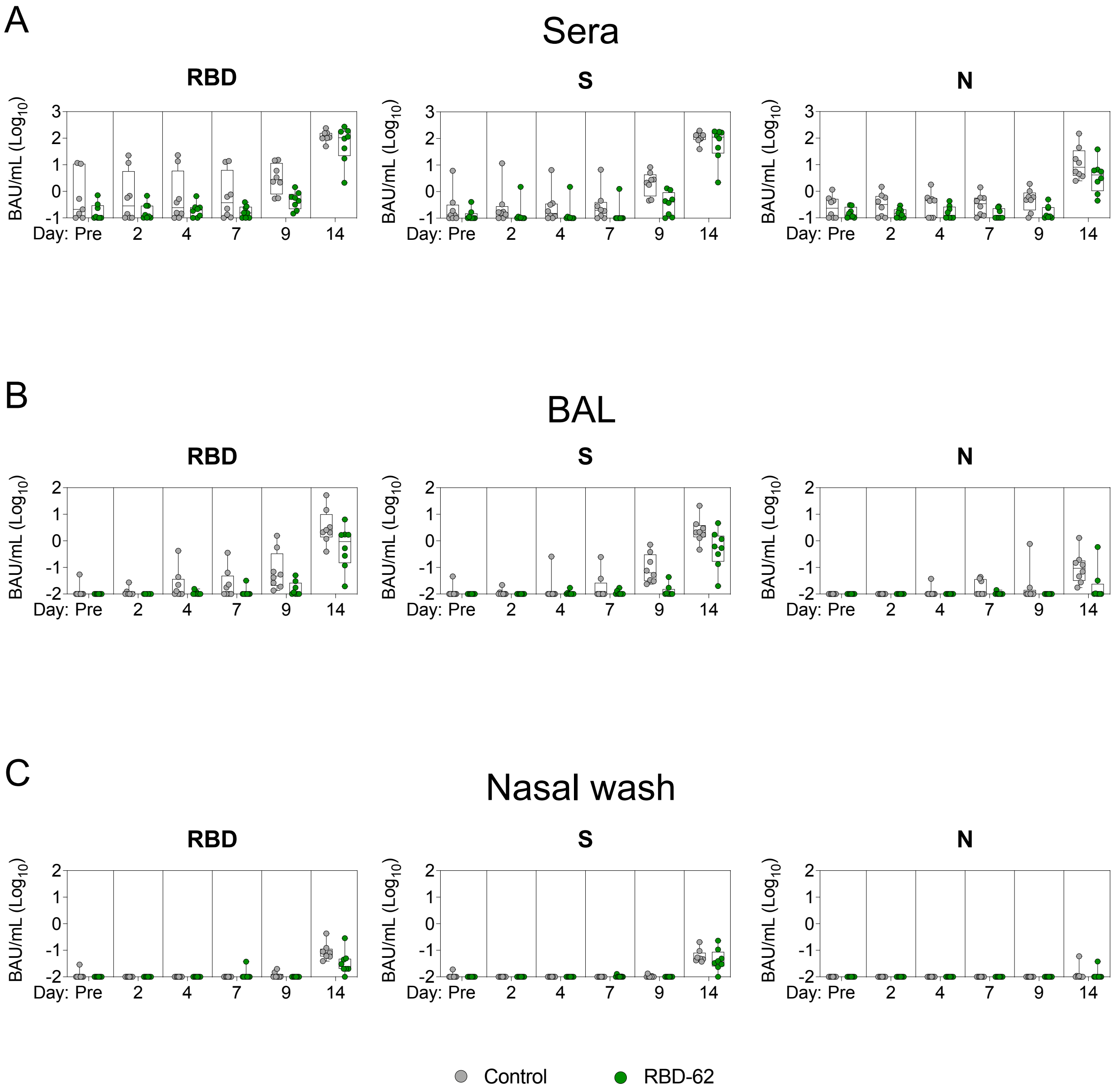


**Fig. S2.**
**Humoral responses to SARS-CoV-2 proteins and domains.** NHP ($n$=8 per group) were challenged with 2x10$^5$ $TCID_{50}$ Delta and simultaneously treated with RBD-62 (green circles) or PBS (gray circles). IgG binding titers were measured to wildtype RBD, S and N in the (**A**) sera, (**B**) BAL and (**C**) nasal wash one month prior to challenge (pre-challenge) and on days 2, 4, 7, 9 and 14 post-challenge. Binding titers in WHO-specified international units may extend beyond the lower range of the y-axis. Sera were diluted at ratios of 1:25, 1:50 and 1:100 while mucosal fluids were diluted at ratios of 1:5, 1:10 and 1:20. Circles, boxes and horizontal lines represent individual animals, interquartile range and median, respectively.

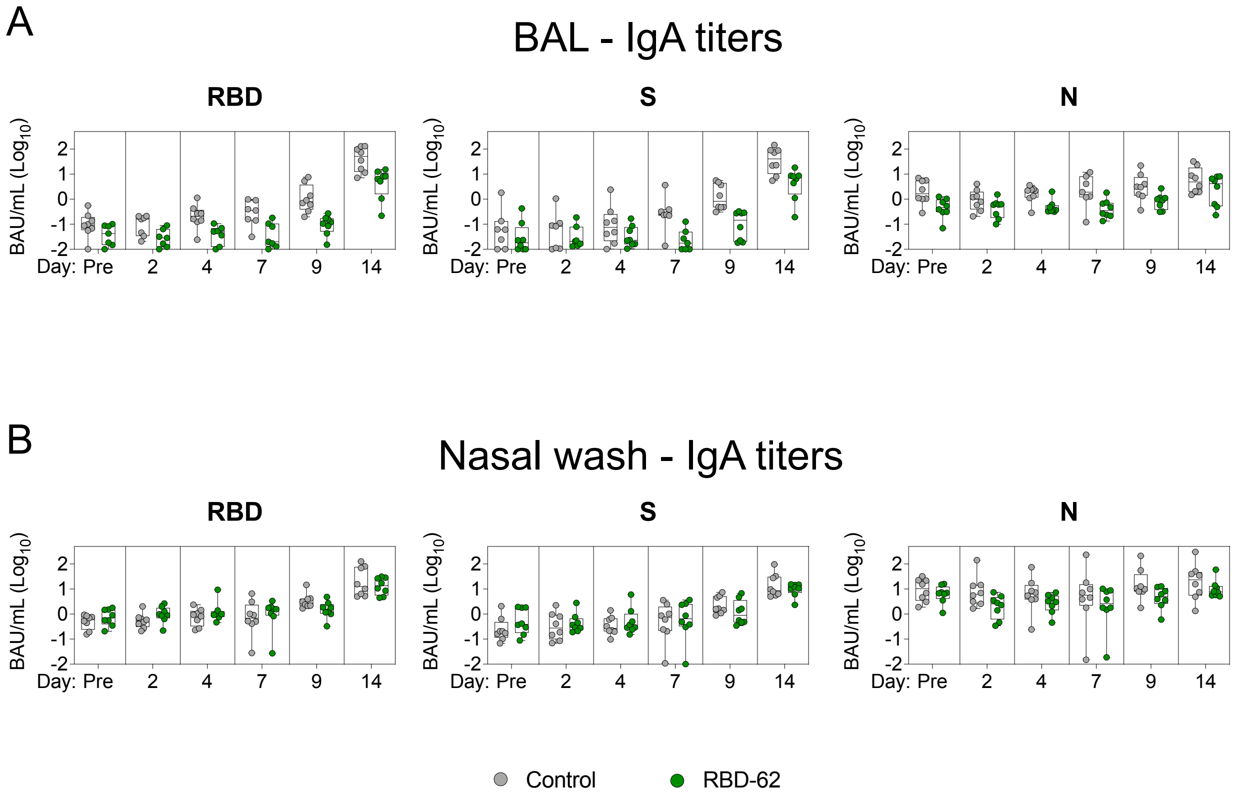


**Fig. S3.**

**Mucosal antibody responses to SARS-CoV-2 proteins and domains.** NHP (*n*=8 per group) were challenged with $2x10^5$ $TCID_{50}$ Delta and simultaneously treated with RBD-62 (green circles) or PBS (gray circles). IgA binding titers were measured to wildtype RBD, S and N in the (**A**) BAL and (**B**) nasal wash one month prior to challenge (pre-challenge) and on days 2, 4, 7, 9 and 14 post-challenge. Binding titers in WHO-specified international units may extend beyond the lower range of the y-axis. Mucosal fluids were initially diluted 1:5 and then serially diluted 1:5. Circles, boxes and horizontal lines represent individual animals, interquartile range and median, respectively.

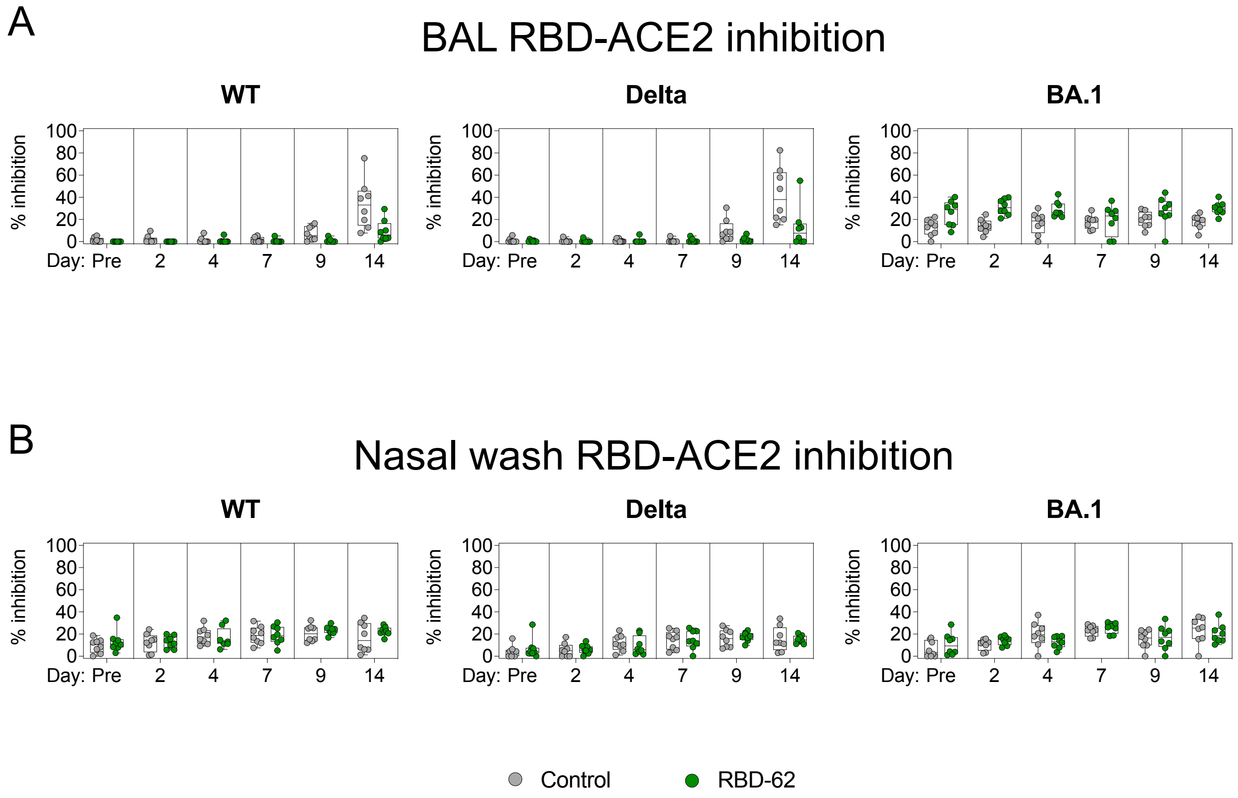


**Fig. S4.**

**RBD-ACE2 binding inhibition in the mucosa.** NHP (*n*=8 per group) were challenged with $2x10^5$ $TCID_{50}$ Delta and simultaneously treated with RBD-62 (green circles) or PBS (gray circles). RBD from WT, Delta and BA.1 variants were mixed with soluble ACE2 in combination with mucosal antibodies from the (**A**) BAL or (**B**) nasal wash one month prior to challenge (pre-challenge) and on days 2, 4, 7, 9 and 14 post-challenge. Percentage binding inhibition was determined relative to maximum binding without inclusion of mucosal fluid. All samples were diluted 1:5. Circles, boxes and horizontal lines represent individual animals, interquartile range and median, respectively.

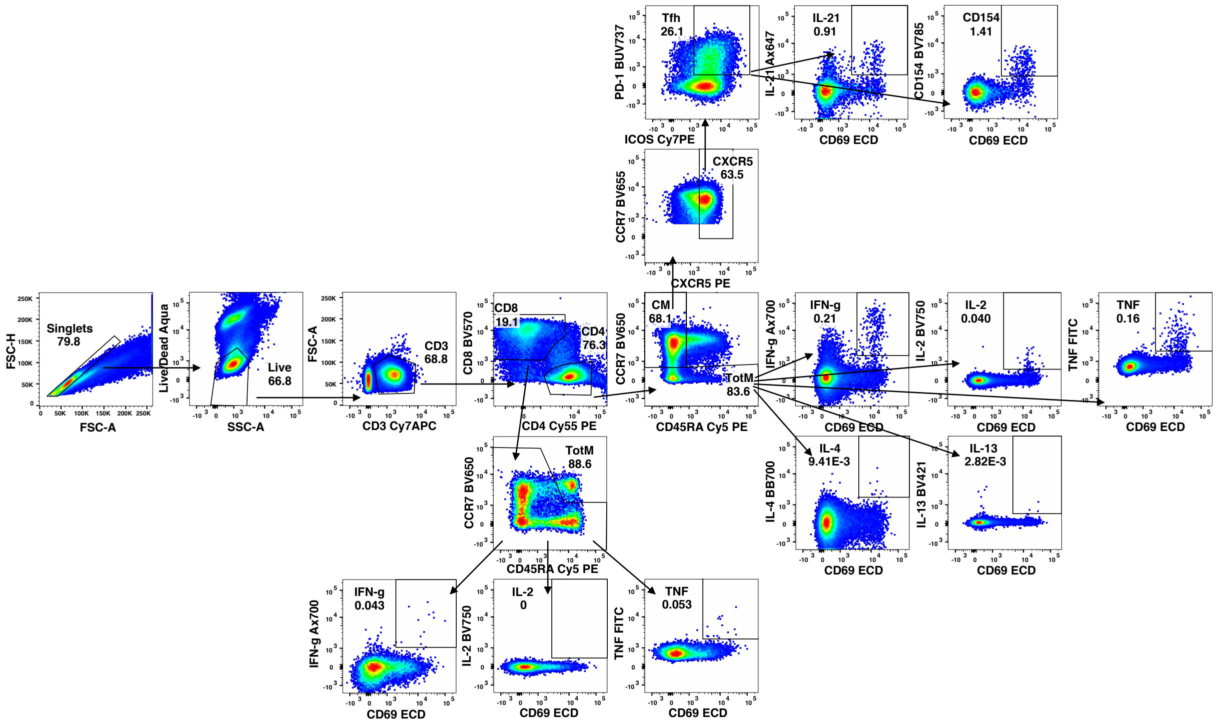


**Fig. S5.**
**ICS gating strategy.** Cells were gated as singlets and live cells on forward and side scatter and a live/dead aqua blue stain. $CD3^+$ events were gated as $CD4^+$ or $CD8^+$ T cells. Total memory $CD8^+$ T cells were selected based on expression of CCR7 and CD45RA. Finally, SARS-CoV-2 S-specific memory $CD8^+$ T cells were gated according to co-expression of CD69 and IL-2, TNF or IFNγ. The $CD4^+$ events were defined as total memory or central memory according to expression of CCR7 and CD45RA. $CD4^+$ cells with a $T_{H}1$ phenotype were defined as memory cells that co-expressed CD69 and IL-2, TNF or IFNγ. $CD4^+$ cells with a $T_{H}2$ phenotype were defined as memory cells that co-expressed CD69 and IL-4 or IL-13. $T_{FH}$ cells were defined as central memory $CD4^+$ T cells that expressed CXCR5, ICOS and PD-1. $T_{FH}$ cells were further characterized as $IL\text{-}21^+$, $CD69^+$ or $CD40L^+$, $CD69^+$.

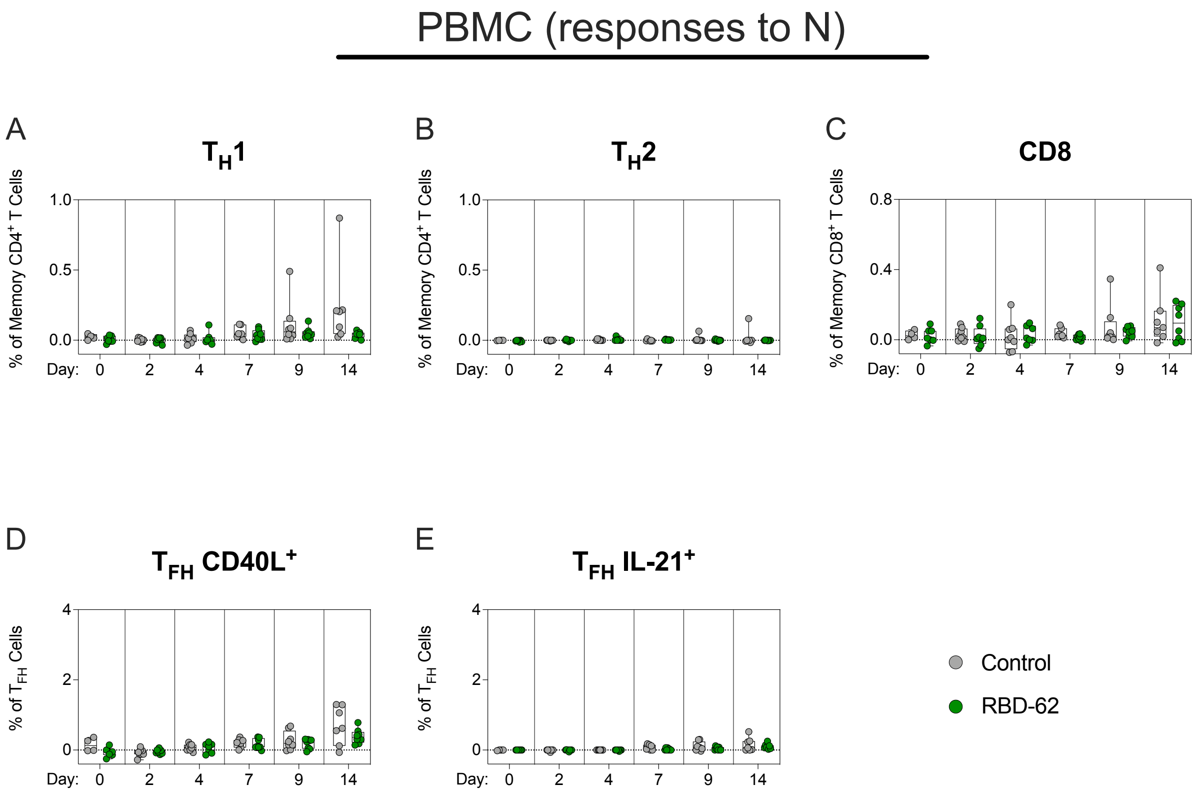


**Fig. S6.**

**Kinetics of primary T cell responses to N peptides following Delta challenge.** NHP (*n*=8 per group) were challenged with $2x10^5$ $TCID_{50}$ Delta and simultaneously treated with RBD-62 (green circles) or PBS (gray circles). Peripheral blood mononuclear cells (PBMC) were collected immediately prior to challenge and on days 2, 4, 7, 9 and 14 post-challenge. Cells were stimulated with WA1 N peptide pools and responses measured by intracellular cytokine staining (ICS). (**A**) Percentage of memory $CD4^+$ T cells expressing $T_H1$ markers (IL-2, TNF or IFNγ). (**B**) Percentage of memory $CD4^+$ T cells expressing $T_H2$ markers (IL-4 or IL-13). (**C**) Percentage of memory $CD8^+$ T cells expressing IL-2, TNF or IFNγ. (**D** and **E**) Percentage of $T_{FH}$ cells expressing CD40L or IL-21, respectively. Dotted lines set at 0%. Reported percentages may be negative due to background subtraction. Circles, boxes and horizontal lines represent individual animals, interquartile range and median, respectively.

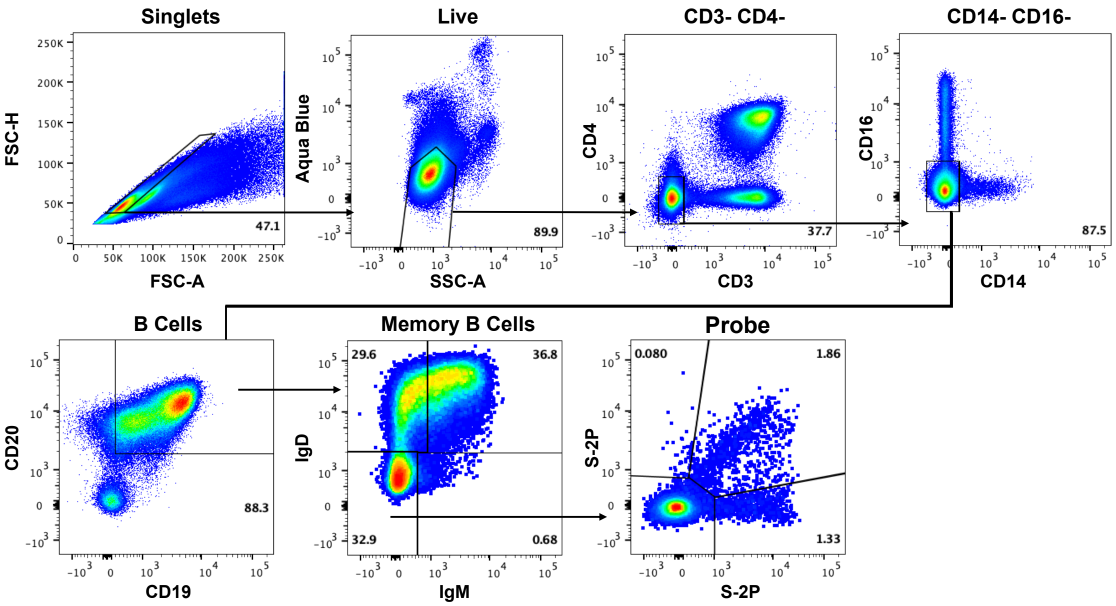


**Fig. S7.**
**Memory B cell gating strategy.** Cells were gated as singlets and live cells on forward and side scatter and a live/dead aqua blue stain. Cells were further gated based on lack of expression of CD3, CD4, CD14 and CD16. B cells were then defined based on expression of CD20 and CD19 whereas memory B cells were gated based on lack of IgD or IgM expression. Finally variant S-2P probe pairs were used to define binding specificity.

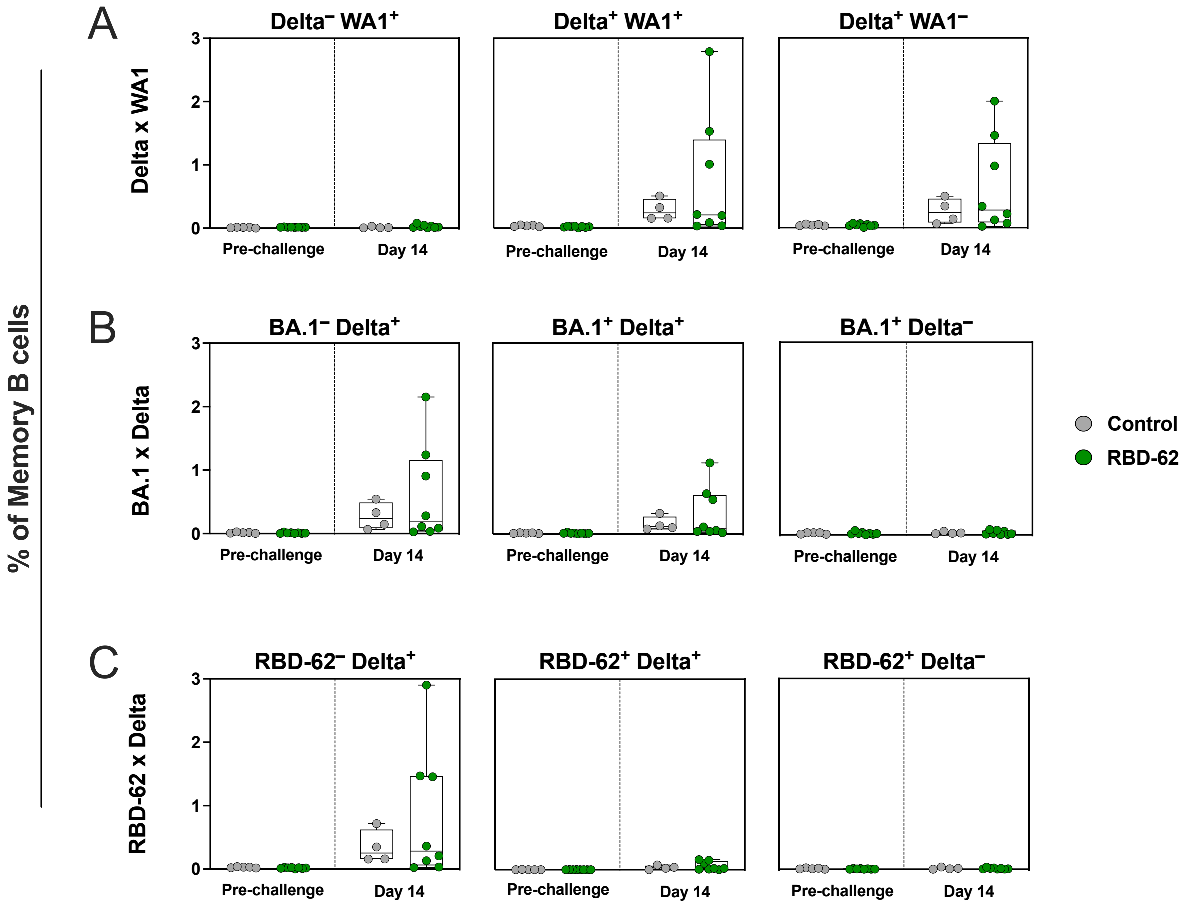


**Fig. S8.**

**Variant-specific and cross-reactive memory B cell frequencies.** NHP (*n*=8 per group) were challenged with 2x10$^5$ TCID$_{50}$ Delta and simultaneously treated with RBD-62 (green circles) or PBS (gray circles). Memory B cells were collected from the periphery and measured via flow cytometry for binding specificity using pairs of fluorescently-labeled variant probes at day 0 and 14 post-challenge. Probe pairs included (**A**) Delta and WA1 S-2P, (**B**) BA.1 and Delta S-2P and (**C**) RBD-62 and Delta S-2P. Cross-reactive cells shown in middle column while variant-specific memory B cells are displayed in left and right columns. Antigen-specific cells reported as a frequency of total memory B cell population. Circles, boxes and horizontal lines represent individual animals, interquartile range and median, respectively.

| RBD-62 Sequence |
| --- |
| TNLCPFGEVFNATRFASVYAWNRKRFSNCVADYSVLYNSASFSTFKCYGVSPTKLN DLCFTNVYADSFVIRGDEVRQIAPGQTGKIADYNYKLPDDFTGCVIAWNSNNLDS KKGGNYNYLYRLFRKSKLKPFERDTSMEIYQAGNTPCNGVKGFNCYFPLQSYGF RPTYGVGYQPYRVVVLSFELLHAPATVCGPKHHHHHH |

**Table S1.**

**RBD-62 Sequence.** Amino acid sequence of His-tagged RBD-62.
